# Gradient organisation of functional connectivity within resting state networks is present from 25 weeks gestation in the human fetal brain

**DOI:** 10.1101/2023.06.26.546607

**Authors:** Jucha Willers Moore, Siân Wilson, Marianne Oldehinkel, Lucilio Cordero-Grande, Alena Uus, Vanessa Kyriakopoulou, Eugene P Duff, Jonathan O’Muircheartaigh, Mary A Rutherford, Laura C Andreae, Joseph V Hajnal, A David Edwards, Christian F Beckmann, Tomoki Arichi, Vyacheslav R Karolis

**Affiliations:** Centre for the Developing Brain, School of Biomedical Engineering and Imaging Sciences, King’s College London, London SE1 7EH, UK; Centre for Developmental Neurobiology, Institute of Psychiatry, Psychology & Neuroscience, King’s College London, London, UK; MRC Centre for Neurodevelopmental Disorders, King’s College London, London, UK; Department of Cognitive Neuroscience, Radboud University Medical Centre, Nijmegen, 6500 HB, The Netherlands; Donders Institute for Brain, Cognition and Behaviour, Radboud University Medical Centre, Nijmegen, 6525 EN, The Netherlands; Biomedical Image Technologies, ETSI Telecomunicación, Universidad Politécnica de Madrid & CIBER-BBN, Madrid, Spain; Biomedical Engineering Department, School of Imaging Sciences and Biomedical Engineering, King’s College London, St. Thomas’ Hospital, London, UK; Department of Brain Sciences, Faculty of Medicine, Imperial College London, UK; UK Dementia Research Institute, Imperial College London, London, W12 0BZ, UK; Department of Forensic and Neurodevelopmental Sciences, Institute of Psychiatry, Psychology & Neuroscience, King’s College London, London SE5 8AF, UK; Paediatric Neurosciences, Evelina London Children’s Hospital, Guy’s and St Thomas’ NHS Foundation Trust, UK; Wellcome Centre for Integrative Neuroimaging, FMRIB, Nuffield Department of Clinical Neurosciences, University of Oxford, UK

**Author notes:** Authors contributed equally (Tomoki Arichi, Vyacheslav R Karolis).

## Abstract

During the third trimester of human gestation, the structure and function of the fetal brain is developing rapidly, laying the foundation for its connectivity framework across the lifespan. During this juncture, resting state functional MRI can be used to identify resting state networks (RSNs) which mature across gestation to resemble canonical RSNs at full term. However, the emergence of finer grain organisation of connectivity within these RSNs in the fetal brain is unknown. Using in-utero resting state fMRI, we performed connectopic mapping analysis to explore the presence of gradients in functional connectivity organisation of 11 cortical RSNs, known as connectopic maps in fetuses aged 25-37 weeks gestation (GW). We hypothesised that, if present, development of connectopic maps would be network specific in the third trimester of gestation, such that this property would be present within the earlier maturing primary sensory and motor networks before those associated with higher association function. In keeping with this, we found smooth connectopic maps in all of the studied RSNs from 25 GW, with the most spatially consistency across gestational age in the primary sensory and motor networks. Voxel-wise permutation testing of the connectopic maps identified local clusters of voxels within networks that significantly covaried with age, specifically in multisensory processing areas, suggesting multisensory processing may be developing during this period. Our analysis shows that functional gradient organisation is already established in the fetal brain and develops throughout gestation, which has strong implications for understanding how cortical organisation subserves the emergence of behaviour in the ensuing period.

## INTRODUCTION

The cortex is a complex and diverse structure that supports a vast array of cognitive processes across the lifespan. It has long been known that there is macroscale regional functional specialisation within the cortex whereby specific functions, such as motor control or visual processing, are localised to specific cortical areas (Glasser et al., 2016; Roland & Zilles, 1998). More recently, it is increasingly appreciated that brain activity occurs in a correlated manner across ensembles of neurons involving different brain regions, known as networks. Synchronised slow fluctuations in cortical activity (functional connectivity) can be identified even in the absence of a task using Blood Oxygen Level Dependent (BOLD) functional MRI (fMRI) as resting state networks (RSNs). Together, RSNs encompass the entire brain, including the primary sensory cortices and higher order cortices (Biswal et al., 1995; Cordes et al., 2000; Damoiseaux et al., 2006; Greicius et al., 2003; Lowe et al., 1998). Importantly, coordinated activity both within and across these RSNs has been found to be especially important for the complex brain processing thought to underlie human behaviour and cognition (van den Heuvel & Hulshoff Pol, 2010).

Historically, neuroimaging has been focused on segmenting the cortex into distinct regions based on functional specialisation whereby one region is responsible for a single function and/or shows a consistent connectivity profile throughout (Han et al., 2018). However, it has since become apparent that functional connectivity is more likely to be topographically organised across the entire brain (rather than parcellated into distinct regions), with connectivity gradually changing continuously across the cortex. This can be characterised as a connectivity gradient such that adjacent locations of the cortex are functionally connected to similar or overlapping cortical areas (Bernhardt et al., 2022; Goldberg, 1989; Haak et al., 2018; Margulies et al., 2016; Sepulcre et al., 2012). This gradient organisation of functional connectivity is now thought to be a fundamental property of the mature human brain which underlies efficient complex neural processing and information integration (Bernhardt et al., 2022; Huntenburg et al., 2018; Kaas, 1997; Laughlin & Sejnowski, 2003; O’Rawe & Leung, 2020). Gradient organisations of functional connectivity within networks, known as connectopic maps, can be used to probe the specificity of connections within a network, for example, the widely recognised topographic organisation of activity and connections within the primary sensory and motor cortices (e.g., somatotopic, tonotopic and retinotopic) (Haak et al., 2018; Park et al., 2021). Alterations in this gradated organisation can be used to probe changes in functional connectivity organisation under different conditions, for example in neurodevelopmental conditions, such as autism (Hong et al., 2019; Park et al., 2021).

Development of the human cortex is a complex, protracted process beginning *in utero* around 8 weeks gestation (GW) and proceeding well beyond birth and childhood (Bystron et al., 2008; Gilmore et al., 2018; Stiles & Jernigan, 2010). Spontaneous cortical activity, which can be studied *in vivo* using fMRI as early as 22 GW (Schöpf et al., 2012), is a hallmark of the developing cortex, emerging during the fetal period and evolving across gestation. Such large scale spontaneous activity is seen to be already organised into RSNs in the third trimester *in utero*, within which functional connectivity increases in complexity and long-range spatial representations become more prominent with increasing gestational age (Karolis et al., 2023; Thomason et al., 2013, 2015; Turk et al., 2019; M. I. van den Heuvel et al., 2018), resulting in RSNs with bilateral and distributed topographies that resemble mature canonical RSNs as identified *ex utero* by the time of normal birth (Doria et al., 2010; Eyre et al., 2021; Fitzgibbon et al., 2020). However, the emergence of finer grain gradient organisation of the underlying connectivity during the fetal period has not yet been explored. Thus, it is unclear if these connectopic gradients are an intrinsically determined property which have a role in shaping the brain’s later cortical networks or whether they emerge based on extrinsic influences. Furthermore, their developmental sequelae may differ across regions and systems, considering that primary sensory and motor RSNs are established early during development, whereas higher order association RSNs are not fully established until early childhood (Doria et al., 2010; Gao et al., 2009; Smyser et al., 2010).

In this study, we aimed to determine the presence of local gradient organisation in functional connectivity of the human fetal brain using *in utero* resting state fMRI data from 151 fetuses aged 25 to 37 GW as part of the developing Human Connectome Project. We performed connectopic mapping analysis (Haak et al., 2018) within 11 resting state networks previously described in neonates (Eyre et al., 2021). We hypothesised that, if present, connectopic gradient development would be network specific in the third trimester, such that it would be present first within the primary sensory and motor networks that are known to mature earlier than those associated with higher association function.

## RESULTS

Fetal resting state fMRI data were acquired in 151 subjects aged 25-37 GW as part of the developing Human Connectome Project (dHCP; www.developing-connectome.org; Fig. 1). All fetal brain images were reviewed and reported as showing appropriate appearances for their gestational age with no evidence of acquired lesions or congenital malformations by an experienced perinatal neuroradiologist. The data were preprocessed using a state-of-the-art fetal specific pipeline including MB-SENSE image reconstruction, distortion correction, slice-to-volume motion correction, temporal denoising, and registration to a weekly template and common template space for group-level analyses (Cordero-Grande et al., 2018; Karolis et al., 2023; Price et al., 2019). Fetal gradients of functional connectivity were characterized using functional connectopic mapping implemented with the Congrads toolbox (Haak et al., 2018). Connectopic mapping aims to characterise how functional connectivity with the rest of the cortex changes across a given region, thus probing the specificity of connections of that individual region. Regions of interest were defined from independently delineated neonatal RSNs (Eyre et al., 2021), to provide a detailed representation of the regional functional specialisation that the fetal cortex is developing towards (Fitzgibbon et al., 2020). Group level connectopic maps were derived for each weekly age group in 11 RSNs in each hemisphere independently to build a trajectory of connectopic maps across gestation in the fetal cortex. We assessed the spatial similarity of derived connectopic maps across gestation and performed a voxel-wise analysis to identify age-related differences in the maps. This allowed us to detect network specific differences in developmental trajectories during a specific period of rapid hierarchical cortical development.

**Figure 1.**
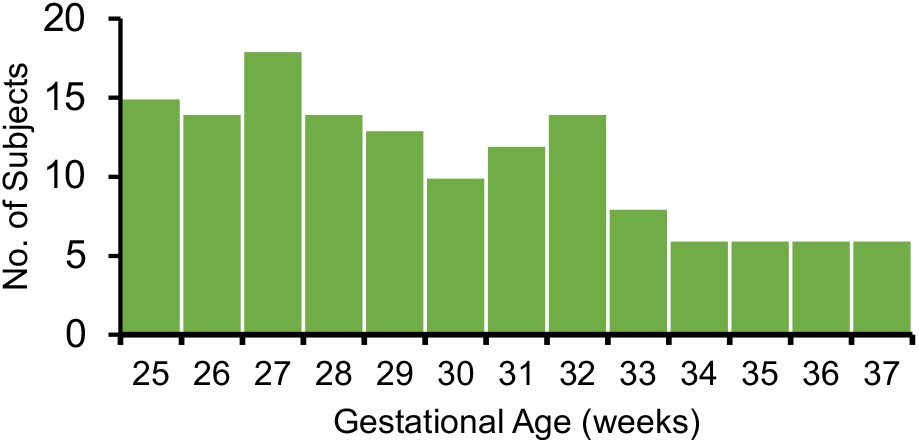
Subject demographics of fetal fMRI datasets.

### Connectopic mapping of primary sensory and motor networks

Connectopic mapping of the primary sensory and motor networks in the fetal brain revealed distinct connectopic gradients from as early as 25 GW in each network (Fig. 2A). The lateral motor and somatosensory networks showed an inferior-superior gradient organisation, the medial motor network showed a medial to lateral gradient and the auditory network showed an anterior to posterior gradient organisation. For the primary visual network, both a principal connectopic gradient with an inferior-superior organisation and a secondary connectopic gradient with an anterior-posterior organisation were identified in the fetal cortex (Fig. 2A), consistent with connectopic mapping work in adults where two distinct gradients representing polar angle and eccentricity of vision are seen (Haak et al., 2018; Watson & Andrews, 2022). Assessing the global spatial similarity of each gradient across gestational age showed that the connectopic maps were remarkably replicable and symmetrical in topography, both across hemispheres and gestation, despite being derived independently (Fig. 2B), as demonstrated by a high mean spatial similarity and a low standard deviation across age.

**Figure 2.**
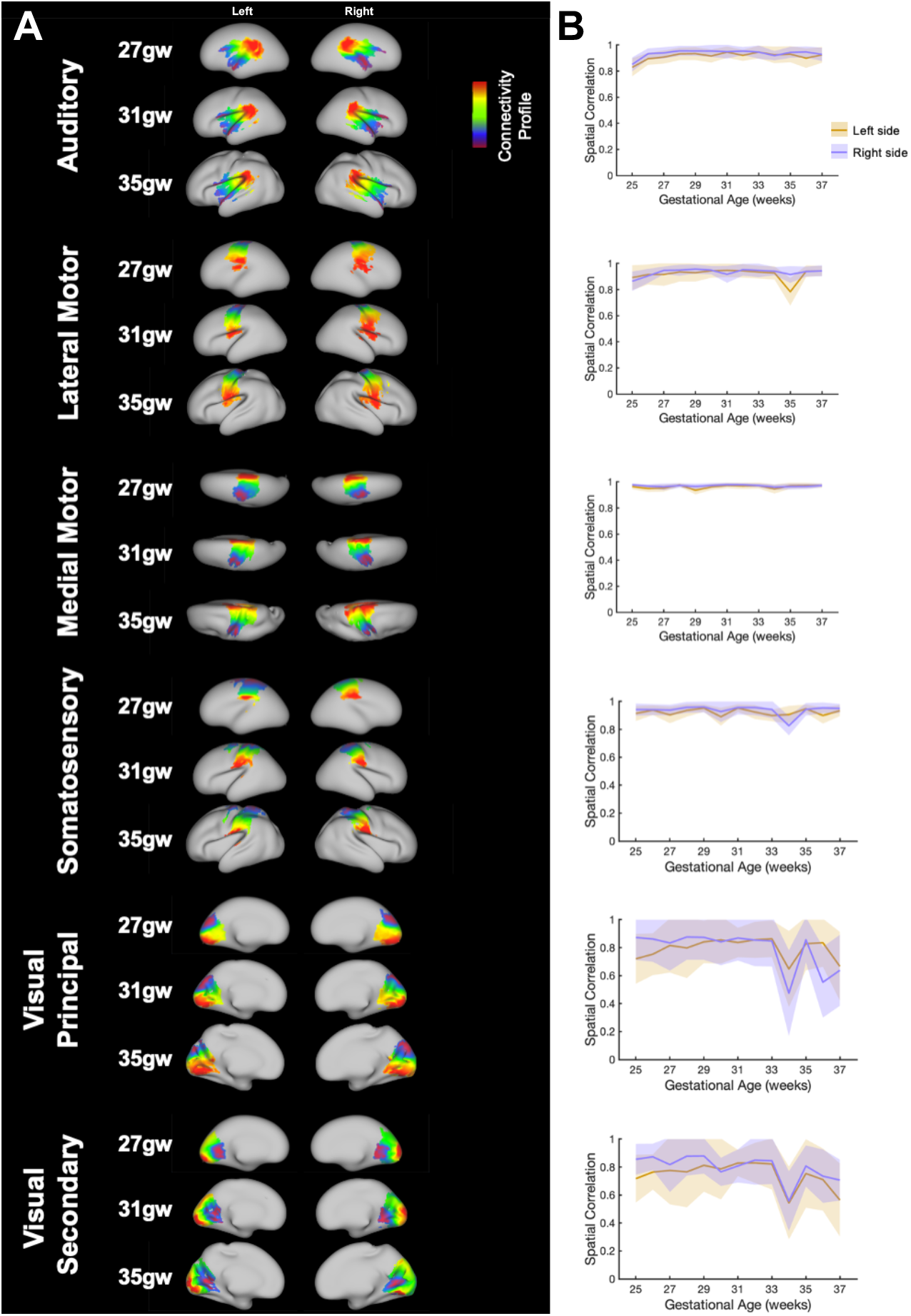
Fetal connectopic mapping analysis in primary sensory and motor resting state networks. (A) Group average connectopic gradient maps of the primary sensory and motor resting state networks in fetuses aged 27, 31 and 35 GW (n = 12, 18 and 6 respectively) overlaid onto high resolution fetal inflated surface atlas images of corresponding gestational age. Similar colours denote similar connectivity profiles. (B) Mean (+/- SD) of the spatial correlation of connectopic gradient maps with all other ages (25-37GW; n=6-18) in the left (orange) and right (purple) hemispheres (right).

Local differences in the connectopic maps were assessed using voxel-wise permutation analysis to identify small clusters of voxels within the networks which positively or negatively significantly co-varied with age, indicative of local changes in functional organisation. Here, the polarity of correlation is inconsequential, it simply denotes a change in the gradient and thus small changes in connectivity profile across age. These clusters were bilateral in the lateral motor, medial motor and secondary visual gradients (Fig. 3A). The auditory and medial motor also showed unilateral voxel clusters which significantly co-varied with age (Fig. 3A). There was no significant change in gradient values across gestation in the primary somatosensory and primary visual gradients.

**Figure 3.**
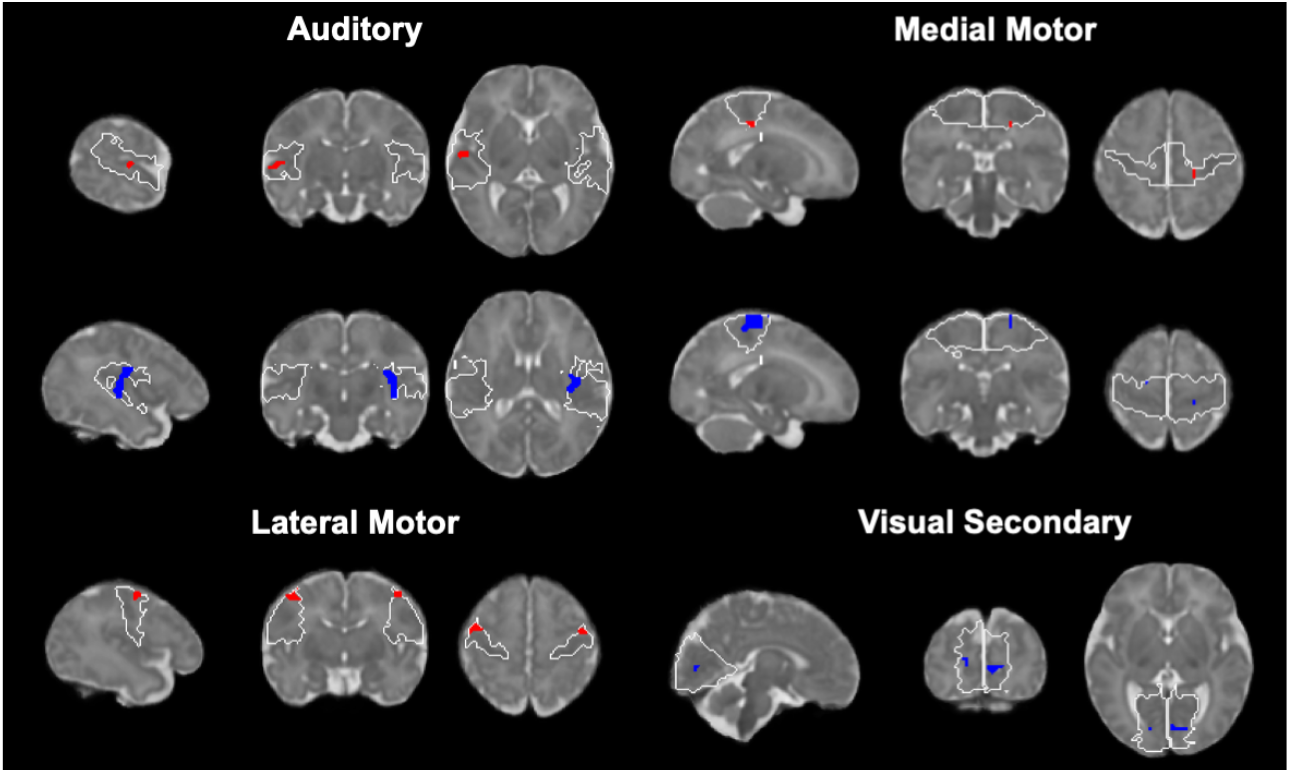
Evolution of fetal connectopic maps with gestational age in primary sensory and motor resting state networks. (A) Voxel clusters within connectopic maps with a significant positive (red) or negative (blue) correlation of gradient values with gestational age for each of the primary sensory and motor networks overlaid onto a high-resolution fetal atlas T2 weighted image at 37 GW (p<0.025).

### Connectopic mapping of visual association and motor association networks

Connectopic mapping of the visual association and motor association networks revealed an inferior-superior gradient in the visual association network and a medial to lateral gradient in the motor association network (Fig. 4A), but with a lower spatial similarity across gestational age (Fig. 4B) as shown by a lower mean spatial similarity and larger standard deviation across age in comparison compared with primary networks. There were large differences in spatial similarity of connectopies across gestation between hemispheres in both these networks (Fig. 4B), with the left hemisphere showing a larger standard deviation of spatial similarity across age compared to the right hemisphere. Large unilateral clusters of voxels with gradient values significantly covarying with age were identified in both the motor and visual association networks in the left and right hemispheres respectively (Fig. 5A).

**Figure 4.**
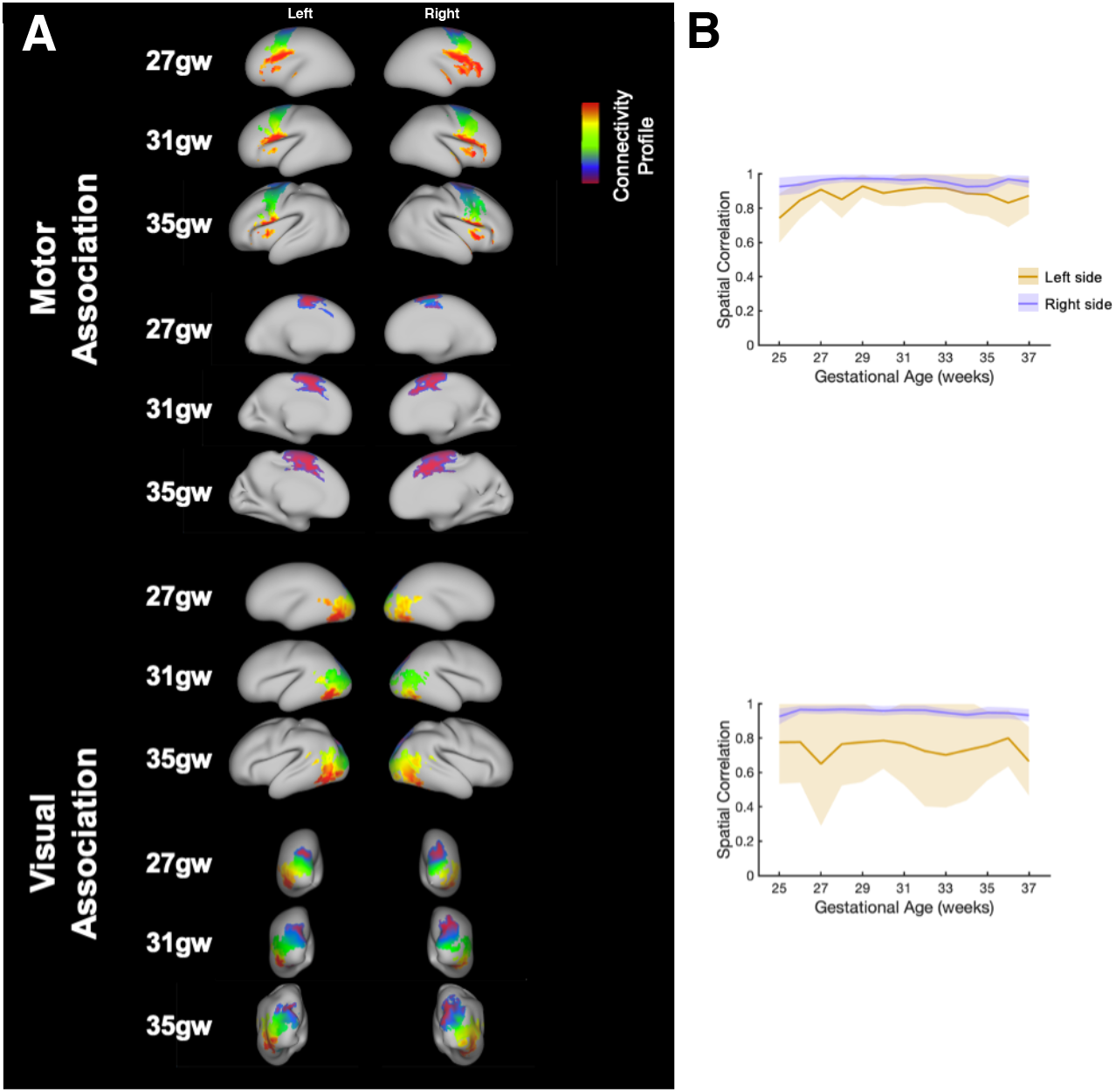
Fetal connectopic mapping analysis in association resting state networks. (A) Group average connectopic gradient maps of the association resting state networks in fetuses aged 27, 31 and 35 GW (n=12, 18 and 6 respectively) overlaid onto high resolution fetal inflated surface atlas images of corresponding gestational age. Similar colours denote similar connectivity profiles. (B) Mean (+/- SD) of the spatial correlation of connectopic gradient maps with all other ages (25-37GW; n=6-18) in the left (orange) and right (purple) hemispheres.

**Figure 5.**
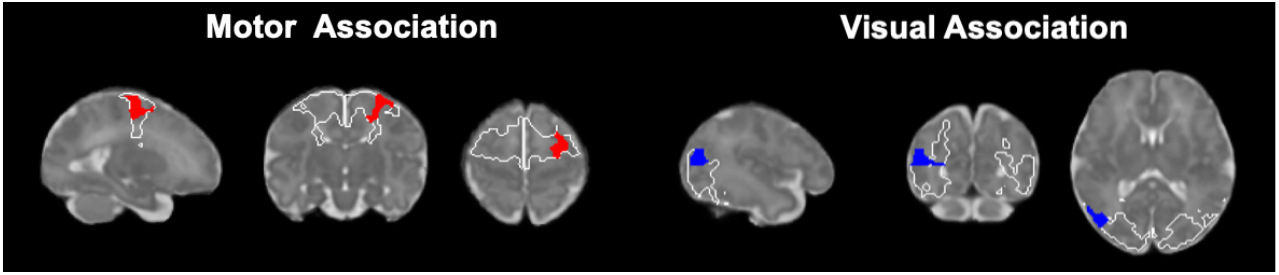
Evolution of fetal connectopic maps with gestational age in association resting state networks. (A) Voxel clusters within connectopic maps with a significant positive (red) or negative (blue) correlation of gradient values with gestational age for each of the association networks overlaid onto a high-resolution fetal atlas T2 weighted image at 37 GW (p<0.025).

### Connectopic mapping of higher-order processing networks

We also performed connectopic mapping of higher-order processing networks, which revealed that functional connectivity within these networks was already organised into connectopic maps early in gestation (Fig. 6A). An anterior to posterior gradient organisation was revealed in both the frontoparietal and temporoparietal networks, whereas an inferior to superior gradient organisation was revealed in the posterior parietal and prefrontal networks. The connectopic maps showed varying degrees of spatial similarity across gestational age, with the temporoparietal network being the most spatially consistent and the prefrontal and posterior parietal networks being the least (Fig. 6B).

**Figure 6.**
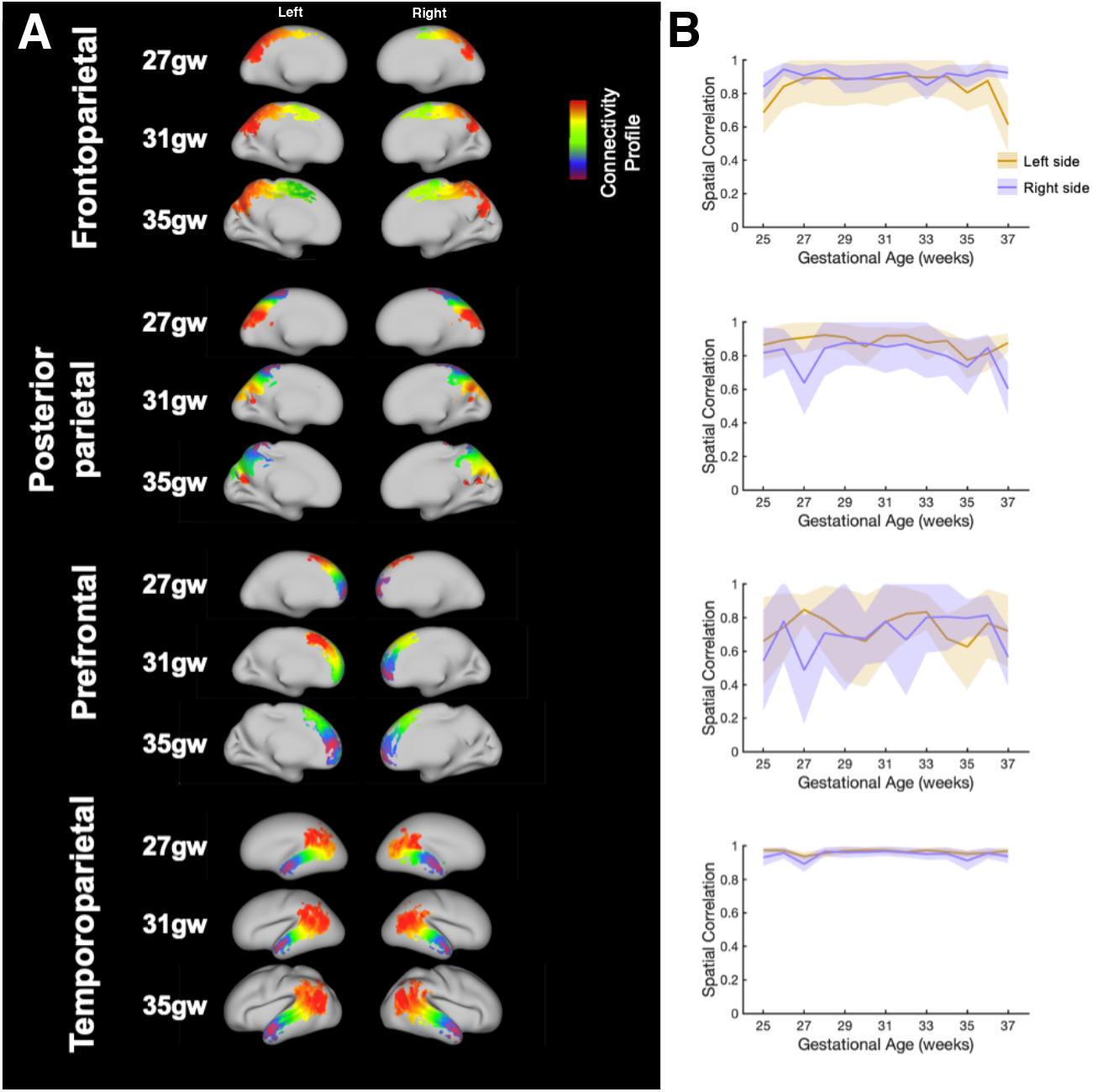
Fetal connectopic mapping analysis in higher-order processing resting state networks. (A) Group average connectopic gradient maps of the higher-order processing resting state networks in fetuses aged 27, 31 and 35 GW (n=12, 18 and 6 respectively) overlaid onto high resolution fetal inflated surface atlas images of corresponding gestational age. Similar colours denote similar connectivity profiles. (B) Mean (+/- SD) of the spatial correlation of connectopic gradient maps with all other ages (25-37GW; n=6-18) in the left (orange) and right (purple) hemispheres.

Voxel-wise analysis of the gradient values across age revealed large bilateral clusters of voxels with gradient values that significantly varied with gestational age in the posterior parietal gradient and unilateral clusters in the frontoparietal and temporoparietal networks in the left and right hemispheres respectively (Fig. 7). No significant voxel-wise changes across age were seen in the prefrontal connectopic map, likely due to the high variability in the topographic organisation of this gradient across gestational age (Fig.6B).

**Figure 7.**
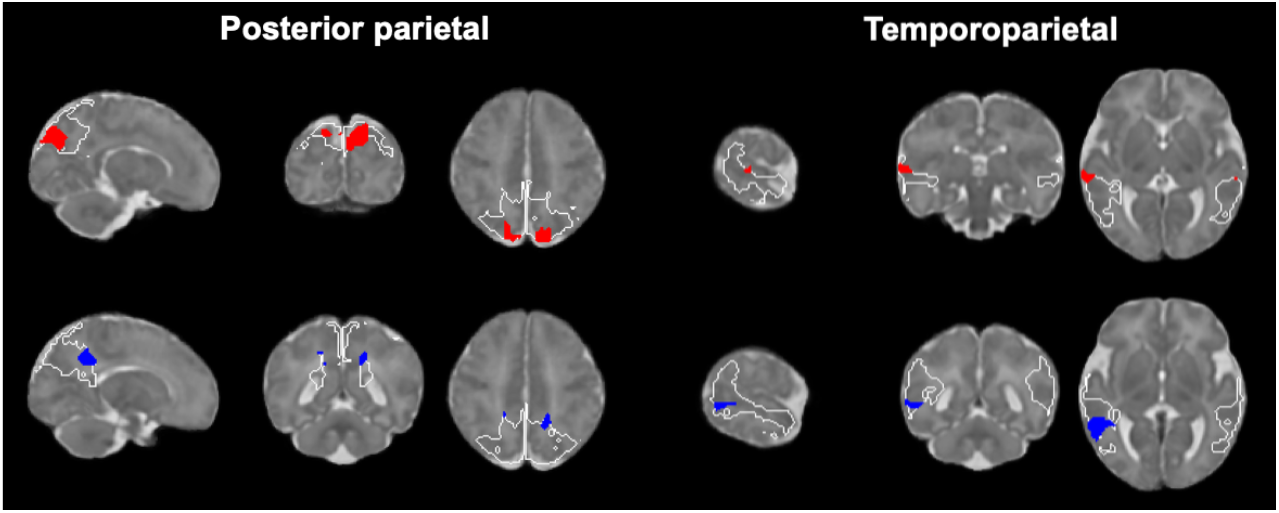
Evolution of fetal connectopic maps with gestational age in higher-order processing resting state networks. (A) Voxel clusters within connectopic maps with a significant positive (red) or negative (blue) correlation of gradient values with gestational age for each of the higher-order processing networks overlaid onto a high-resolution fetal atlas T2 weighted image at 37 GW (p<0.025).

## DISCUSSION

Gradated organisation of functional connectivity is thought to be integral to efficient information processing and transfer via local processing streams across the brain (Laughlin & Sejnowski, 2003; Margulies et al., 2016; O’Rawe & Leung, 2020). In this study, we show that robust gradient organisations of functional connectivity are already established within the majority of RSNs in the fetal cortex from 25 GW. Globally, this organisation remains largely stable across the studied age range, suggesting that the organisation of these RSNs is shaped early in gestation and is intrinsically determined. This was most evident in the primary sensory and motor networks, which showed remarkably consistent connectopies across our study period of 25-37 GW that are already comparable to the canonical topographic organisation of functional connectivity in corresponding cortical regions in the mature adult brain (Haak et al., 2018; Ngo et al., 2021; Park et al., 2021; Watson & Andrews, 2022). Within this, we have identified local clusters of age-related differences in the gradient values, suggesting an on-going refinement of their connectivity structure. Surprisingly, even though higher order and associative networks are the last to be established during development (Gilmore et al., 2018), we identified a gradient organisation of functional connectivity within these networks even in the fetal period. However, gradients in these networks are comparatively less robust, showing greater changes across gestational age, in line with the regional heterochronicity of cortical development (Cadwell et al., 2019).

Studies in neonates suggest that RSN organisation emerges during a period of rapid neural growth and establishment of thalamocortical projections (Dall’Orso et al., 2022; Doria et al., 2010; Eyre et al., 2021). Even though RSNs emerge prior to acquiring the cognitive competencies which are associated with them, it is not clear whether finer grain functional organisation of the networks is also present during this process of development, or emerges in an experience dependent manner. In the case of the primary somatosensory and motor cortices, somatotopic activation maps and patterns of connectivity emerge as early as mid-third trimester in preterm neonates (Allievi et al., 2016; Dall’Orso et al., 2018, 2022). Topographic mapping in the rodent primary sensory cortex similarly emerges in the equivalent of the third trimester and is driven by genetically determined cues and incoming spontaneous peripheral activity (Seelke et al., 2012). Our results demonstrate that in the human fetal cortex, finer grain organisations of RSNs are indeed present from 25 GW. Furthermore, the topographies of the fetal functional gradients we have identified here are remarkably reminiscent of the functional organisation present in the mature lateral and medial motor gradients, principal and secondary visual gradients, and somatosensory gradient (Haak et al., 2018; Ngo et al., 2021; Park et al., 2021; Watson & Andrews, 2022).

Regional functional specialisation within the cortex is tightly constrained by its underlying cytoarchitecture and structural connectivity (Passingham et al., 2002). Traditionally, there have been two main schools of thought regarding how regional specialisation emerges in the developing cortex. The first (“tabula rasa”) proposes a homogenous cortex where any region has the potential to assume every function, with regional specialisation defined by afferent inputs and environmental influence through activity dependent mechanisms (Creutzfeldt, 1977). The alternative hypothesis proposes that regional functional and cytoarchitectural specialisation is predetermined by a cortical protomap implemented through genetics and molecular signalling mechanisms, with later refinement by thalamic pathways (“the radial-unit hypothesis”) (Rakic, 1988). In line with the radial-unit hypothesis, we revealed that the topography of network functional organisation in most gradients remained largely consistent from 25-37 GW in the fetal brain, suggesting it is minimally shaped by *ex utero* influences in this period.

Despite the overall topography of network functional connectivity remaining consistent across gestational age, we demonstrated that there are small areas within networks in which gradient values are changing with increasing gestational age, which could indicate local changes in functional organisation. Localisation of these voxel clusters by comparison with a regional fetal atlas (Gholipour et al., 2017) suggests involvement in multimodal sensory integration and processing in the mature brain, for example, the superior temporal gyrus (Calvert, 2001) in the auditory and temporoparietal networks, the precuneus (Cavanna & Trimble, 2006) in the posterior parietal network, and the middle occipital gyrus (Beauchamp, 2005) in the visual association area. As cortico-cortical and thalamocortical connections are also required for multimodal integration of sensory information, thalamocortical function is likely developing concomitantly with connectivity from 26-34 GW, commissural connectivity from 28-34 GW and cortical associative connectivity from 35-37 GW (Keunen et al., 2017; Kostović et al., 2021). In keeping with this, increased integration of sensorimotor associative areas such as the supplementary motor area and ipsilateral peri-rolandic regions is seen in sensorimotor functional responses with increasing age in preterm infants aged 30-43 weeks post menstrual age (PMA) (Allievi et al., 2016; Whitehead et al., 2019). Recent work has demonstrated that the developing brain can learn associations between multisensory inputs by altering activity in the sensory cortices from 38 weeks PMA (Dall’Orso et al., 2021). It has further been demonstrated that some aspects of multisensory perception are present from 1 month of age, with increasing functionalities of multisensory perception emerging with increasing age into infanthood (Burr & Gori, 2012). Here, our results suggest that the connectivity architecture required for multimodal sensory processing may begin emerging within the window captured by our study period, prior to 38 weeks PMA.

Regional specialisation of cortical function can be situated along a hierarchical axis from primary sensory and motor processing to higher level processing of executive function in networks such as prefrontal and default mode networks (Mesulam, 1998). In the fetal period and through to early childhood, cortical structural and functional development is thought to progress along this same primary to associative axis (Gilmore et al., 2007, 2018; Gogtay et al., 2004). In agreement with this, neonates show an adult like configuration of the primary sensory network by 37 weeks PMA with no significant changes in the architecture to term age (Eyre et al., 2021). Higher order association networks, on the other hand, are still developing from 37-43 weeks PMA (Eyre et al., 2021). Our connectopic mapping analysis also demonstrated a highly consistent topographic organisation within the primary sensory and motor networks and crucially, also found less robust connectopic maps with lower spatial consistency across age in the association and higher processing networks. A key example is seen in the early posterior parietal network, a likely precursor of the default mode network (DMN), which exhibits an adult-like organisation by 2 years of age (Gao et al., 2009). Here we identified large regions of the posterior parietal network which changed significantly with age. This alludes towards the rudimentary structural and functional framework of the brain being largely established even before the development of the cognitive competencies associated with stimulus independent thought (Doria et al., 2010; Gilmore et al., 2018).

## CONCLUSION

Cortical development in the human fetal brain is a complex and protracted process with multiple overlapping developmental events. We demonstrate that a gradient organisation of functional connectivity in RSNs is already present in the fetal cortex from 25 GW. This organisation is most striking and robust in the primary sensory and motor networks, indicating that the organisation of functional connectivity is largely intrinsically established early in gestation. Conversely, gradient organisation within association and higher order RSNs showed lower spatial consistency and more extensive changes across development. This highlights regionally dependent developmental trajectories of functional connectivity along an axis from primary sensory processing to higher order processing. This has strong implications for understanding how cortical organisation subserves the emergence of behaviour in the ensuing period. Not only this, disruptions in cortical gradients may ultimately provide valuable information about how cortical processing is altered in disorders emerging in this critical window for neurodevelopment.

## METHODS

### Participants

Fetal fMRI data were collected in 151 fetuses (25.0-37.8 weeks gestation, GW, mean: 31.6 GW) as part of the Developing Human Connectome Project (dHCP) (www.developing-connectome.org).

### Acquisition

fMRI data were acquired with a Philips Achieva 3T system (Best, NL) and a 32-channel cardiac coil using a single-shot echo planar imaging (EPI) (TR/TE = 2200/60ms) sequence consisting of 350 volumes of 48 slices each, isotropic resolution = 2.2 mm, multiband factor = 3, SENSE factor 1.4 (Price et al., 2019). All fetal brain images were reported by a neuroradiologist as having an appropriate appearance for their gestational age and without clinically significant lesions or congenital malformations. fMRI data from 9 fetuses were discarded upon visual inspection due to excessive motion during acquisition or failed image reconstruction.

### Preprocessing

FMRI data were pre-processed using a state-of-the-art fetal-specific pre-processing pipeline (Cordero-Grande et al., 2018; Karolis et al., 2023; Price et al., 2019). Briefly, pre-processing comprised MB-SENSE image reconstruction, dynamic shot-by-shot B0 field distortion correction, slice-to-volume motion correction and temporal denoising using a model combining volume censoring regressors based on signal outliers, highpass (1/150Hz) filtering regressors, 6 white matter and CSF time courses (obtained using subject-level ICA within a combined white matter + CSF mask) (Behzadi et al., 2007) and 3 types of voxelwise 4D denoising maps characterising 1) time courses of voxels linked in multiband acquisition to voxels in the original data, 2) temporal evolution of a shrinkage/expansion operator applied to the data in order to compensate for the distortion-related volume alterations and aiming to remove the residual effects of distortion corrections on the data timecourses and 3) the motion parameters, estimated during slice-to-volume reconstruction and expanded to include first- and second-order volume-to-volume and slice-to-slice differentials and their square terms (Karolis et al., 2023; Price et al., 2019; Pruim et al., 2015a; Pruim et al., 2015b). No lowpass temporal filtering was applied.

### Registration to weekly template space

Individual subjects’ functional data were registered from native space to a weekly structural template space (available at https://brain-development.org/brain-atlases/fetal-brain-atlases/) (Serag et al., 2012), by firstly rigidly aligning mean functional images to the subjects’ own anatomical T2 weighted image using FSL FLIRT (Jenkinson & Smith, 2001), then non-linearly transforming data from structural space to age-matched weekly structural template space using ANTS (Avants et al., 2011).

### Registration to common template space

For comparison across ages, functional data were non-linearly aligned to the template space corresponding to the gestational age of the oldest subjects, 37 GW, as this age has the best effective resolution and most complex structural features. This required the concatenation of several intermediate transformations to avoid multiple interpolations, including registration to the weekly template space (as outlined above) as well as a sequence of non-linear transformations between contiguous weekly templates spaces to 37 GW template space (Fitzgibbon et al., 2020; Karolis et al., 2023).

### Functional connectivity gradient mapping

#### Regions of interest

As the spatial distribution of functional networks are known to evolve across gestation, to enable appropriate comparison between ages we used neonatal resting state networks (RSNs) derived from an independent group in the dHCP neonatal dataset as regions of interest (Eyre et al., 2021). The RSNs were thresholded (z-score>3), binarised, registered to the same fetal 37 GW template, downsampled to 2 mm isotropic resolution, and manually divided at the midline to generate left hemispheric and right hemispheric component masks of each network. Networks were mapped from a 40 week PMA neonatal template space to the 37 GW fetal template space using a concatenated sequence of non-linear transformations. This included sequential non-linear transformations between neonatal templates of adjacent weekly ages from 40 weeks PMA to 37 weeks PMA, followed by a nonlinear transformation from neonatal 37 weeks PMA to fetal 37 GW template spaces. Similarly to the registration of the functional data to the fetal template space, the transformations were merged into a single, concatenated transformation to bring the neonatal network masks into the fetal template space with only a single interpolation (Fitzgibbon et al., 2020). A probabilistic cortical grey matter template mask (Serag et al., 2012) was also thresholded at 0.5 and downsampled to 2mm isotropic resolution.

#### Connectopic mapping analysis

Functional connectivity topography was mapped in each of the 11 network ROIs using the Congrads toolbox, as described in depth in (Haak et al., 2018). Connectopic mapping aims to estimate the connectivity between each voxel in an ROI with all other cortical grey matter voxels. In brief, the voxelwise fMRI time-series from each ROI and all other cortical voxels were arranged in time-by-voxels matrices. The voxelwise time series matrix of the cortical voxels underwent dimensionality reduction using singular value decomposition (SVD). The voxelwise time-series matrices from the ROI and the cortex were correlated to generate voxelwise connectivity fingerprints for each voxel within the ROI. We then computed the η2 coefficient, which represents the amount of variation in one connectivity profile that can be explained by the variance in another connectivity profile, where higher values indicate greater similarity in the connectivity profiles. The η2 coefficient was then used to produce a similarity matrix, **S**, characterising the within ROI similarity in voxelwise connectivity fingerprints between the ROI voxels and the cortical voxels in the rest of the brain. **S** was constructed for each subject in a weekly age group (13 weekly age groups, 25-37 GW) and then averaged across subjects. A Laplacian eigenmap non-linear manifold learning algorithm was applied to the averaged **S**, to define multiple overlapping but independent modes of functional connectivity within each ROI. Functional connectivity topographies were mapped, known as connectopic mapping in each network ROI at the group level in 13 weekly age groups (25-37 GW), with independent mapping in each hemisphere and age group. Spatially consistent connectopic maps were selected for each week, regardless of order, however, in the majority of cases, this was gradient 1.

Functional connectopic mapping was carried out in weekly template space, and then again once the subjects’ data had been registered to the common 37 GW template space in order to compare functional connectivity topographies across ages.

### Statistical analysis

#### Global spatial similarity

In order to assess the global similarity of connectopic maps across age (25-37 GW) in each network, we calculated spatial correlation matrices of the connectopic maps in each hemisphere across age using AFNI (Cox, 1996; Cox & Hyde, 1997). The mean and standard deviation were calculated across each row of the spatial correlation matrices to give a measure of global consistency of the connectopic maps at each weekly timepoint.

#### Voxel wise age-related changes

To determine whether the gradients were developing across gestation, voxelwise permutation testing (family-wise error corrections for multiple comparisons) was used to identify voxels within the connectopic maps with gradient values significantly covarying with age (two-tailed, p<0.025), implemented in FSL Randomise v1.0 (Winkler et al., 2014).

## ACKNOWLEDGEMENTS

We thank the patients who agreed to participate in this work and the staff of St. Thomas’ Hospital London. This work was supported by the European Research Council under the European Union Seventh Framework Programme [(FP/2007-2013)]/ERC Grant Agreement No. 319456. We also acknowledge grant support in part from the Wellcome Engineering and Physical Sciences Research Council (EPSRC) Centre for Medical Engineering at King’s College London [WT 203148/Z/16/Z] and the Medical Research Council (UK) [MR/K006355/1] and [MR/L011530/1]. J.W.M and S.W. were supported by PhD funding from the UK Medical Research Council [MR/P502108/1]. J.O. is supported by a Sir Henry Dale Fellowship jointly funded by the Wellcome Trust and the Royal Society [206675/Z/17/Z]. J.O., M.A.R., L.A., A.D.E., J.O. and T.A. received support from the Medical Research Council Centre for Neurodevelopmental Disorders, King’s College London [MR/N026063/1]. L.C-G. received support from Project PID2021– 129022OA-I00 funded by MCIN / AEI / 10.13039/501100011033 / FEDER, EU. T.A. was supported by an MRC Clinician Scientist Fellowship [MR/P008712/1]. V.K. and T.A. were supported by an MRC Transition Support Award [MR/V036874/1]. Support for this work was also provided by the NIHR-BRC at Kings College London, Guy’s and St Thomas’ NHS Foundation Trust in partnership with King’s College London, and King’s College Hospital NHS Foundation Trust.

